# Extinction vortices are driven more by a shortage of beneficial mutations than by deleterious mutation accumulation

**DOI:** 10.1101/2024.10.25.620329

**Authors:** Walid Mawass, Joseph Matheson, Ulises Hernández, Jeremy J. Berg, Joanna Masel

## Abstract

Habitat loss contributes to extinction risk in multiple ways. Genetically, small populations can face an “extinction vortex” — a positive feedback loop between declining fitness and declining population size. Two distinct genetic mechanisms can drive a long-term extinction vortex: i) ineffective selection in small populations allows deleterious mutations to fix, driving “mutational meltdown”, and ii) smaller populations generate fewer beneficial mutations essential for long-term adaptation, a mechanism we term “mutational drought”. To determine their relative importance, we ask whether, for a population near its critical size for persistence, changes in population size have a larger effect on the beneficial vs. deleterious component of fitness flux. In stable environments, we find that mutational drought is nearly as significant as mutational meltdown. Drought is more important than meltdown when populations must also adapt to a changing environment, unless the beneficial mutation rate is extremely high. Linkage disequilibria from background selection under realistically high deleterious mutation rates modestly increase the importance of mutational drought. Long-term conservation efforts should consider adaptive potential, not just deleterious load.

## Introduction

To what degree the genetics of a population might contribute to population decline and eventual demise, and how, remains an open question [1–7]. “Extinction vortex” is an umbrella term for a variety of self-reinforcing feedback loops between population size and fitness, including some based on genetics [8]. On short timescales, a population size reduction will increase inbreeding load via deleterious recessive alleles, lowering fitness and potentially reducing population size further [9,10]. On slightly longer timescales, the loss of standing genetic variation can undermine adaptive potential. Here we focus on two even longer-term kinds of extinction vortex, operating over the timescale of fixation events (figure 1). First, ineffective selection in the face of stronger random genetic drift in small populations might allow a series of deleterious mutations to become fixed, in a process of “mutational meltdown”. Second, fewer individuals entail fewer opportunities for the appearance of the novel mutations required for long-term adaptation, a process we call “mutational drought”.

**Figure 1.**
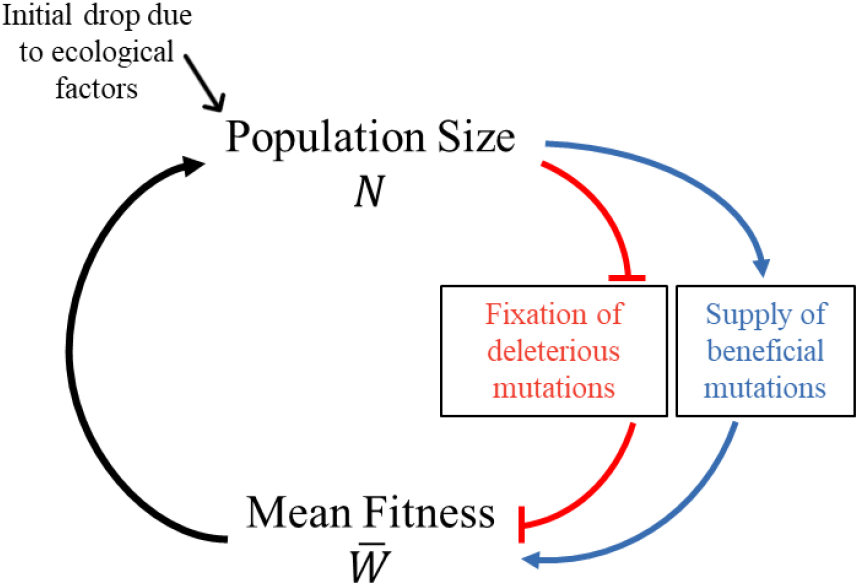
An extinction vortex can be driven by either deleterious (mutational meltdown; red) or beneficial (mutational drought; blue) fixations. When the net effects of deleterious fixations, environmental change, and beneficial fixations cause mean population fitness to decline, the population size *N* will fall. This fall in *N* can further exacerbate the tendency for fitness to fall, creating a positive feedback loop (extinction vortex).

Mutational meltdown has received the most theoretical attention of the two [11–19]. Mutational meltdown is a positive feedback loop between population size *N* and mean fitness 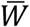. As a population declines in size, deleterious mutations fix more easily due to random genetic drift (Figure 1, top red blunt arrow ┴). This decreases mean fitness 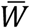 (Figure 1, bottom red blunt arrow), causing the population size to decline further (Figure 1, upward arrow).

Without beneficial mutations, even large populations melt down eventually. This is because some deleterious mutations are too small to be reliably purged by selection, even in large sexual populations; this has been called “Ohta’s ratchet” [20]. With a finite number of sites, back mutations could eventually halt this decline, if we ignore essentially irreversible deletions. But for vertebrate parameter values, this comes at the cost of ∼100 “lethal equivalents” in the exponent of fitness relative to an individual without deleterious mutations [21]. One proposed solution to this problem [22] unrealistically assumes that unconditionally deleterious mutations do not exist, only mutations to quantitative traits under stabilizing selection.

A more compelling solution is to include larger-effect beneficial mutations, not just reversals [23,24]. This enables compensatory asymmetry, where each larger-effect beneficial fixation offsets the fitness effects of multiple small effect deleterious fixations [24–26]. The effects of deleterious and beneficial fixations on population mean fitness can be captured by their respective “fitness fluxes”: 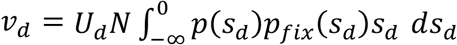 and 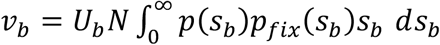, where *U*_*d*_ and *U*_*b*_ give the genome-wide rates of deleterious and beneficial new mutations in each of the *N* individuals, with selection coefficients *s*_*d*_ or *s*_*b*_, respectively, according to probability density function *p*(*s*). Mutations fix with probability *p*_*fix*_(*s*), changing population fitness by *s*.

Schultz and Lynch [23] emphasized the critical ratio of mutation rates *U*_*b*_/*U*_*d*_ that is needed to balance deleterious and beneficial fluxes, as a function of population size *N* and degree of outcrossing. Here we follow Whitlock [24], who studied an essentially identical model, but instead emphasized a critical population size *N*_*crit*_ as a function of *U*_*b*_/*U*_*d*_. Below *N*_*crit*_, a population with that value of *U*_*b*_/*U*_*d*_ crosses the threshold into an extinction vortex.

New beneficial mutations create a second, distinct positive feedback loop (Figure 1, blue arrows). Declining *N* reduces the number of new beneficial mutations, *NU*_*b*_. (See [27] for empirical support that the beneficial mutation rate can be limiting.) This slows the adaptive increase in 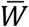, which feeds back into lower *N* (Figure 1, upward arrow). The importance of this second feedback loop was previously briefly noted in the Discussion of [28] as an incidental finding, within a study focused on the effect of fitness function shape on extinction vortices in the context of a rapidly changing environment. We refer to this novel type of extinction vortex as “mutational drought.” Adaptive fixations could be in short supply not only to counter environmental change, but also just to offset deleterious fixations under high *U*_*d*_, even in the absence of environmental change.

Here we ask how important mutational drought is, relative to mutational meltdown. To capture the dynamics of either kind of extinction vortex, fitness must affect population density. In the simplest models of absolute fitness, this leads to unbounded exponential growth for *N* above a threshold *N*_*crit*_, and so many previous meltdown models study dynamics only once *N* < *N*_*crit*_ [12,13,29]. Here we ignore dynamics and instead take a perturbation approach around the tipping point *N*_*crit*_ (which is an unstable equilibrium in phase space), to calculate the derivatives of the two fitness fluxes with respect to population size. A “drought : meltdown ratio” 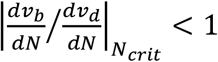 indicates that mutational meltdown is more important in the vicinity of the tipping point, while a ratio > 1 indicates that mutational drought is more important. This perturbation approach allows us to ignore dynamical trajectories. It even allows us to use standard relative fitness models of constant *N*, under the assumption that first order effects of *N* on fitness fluxes are larger than second order feedbacks.

Previous models [23,24] of fitness flux *UN* ∫ *p*(*s*)*p*_*fix*_(*s*)*s ds* assumed in their calculations of *p*_*fix*_(*s*) that each mutation evolves independently. However, the independence assumption is violated, even in sexual populations, by background selection caused by deleterious mutations, and by hitchhiking and clonal interference caused by beneficial mutations [30,31]. These phenomena increase the fixation probabilities of deleterious mutations and decrease those of beneficial mutations.

Altered fixation probabilities for both deleterious [32] and beneficial [33] mutations correspond to linkage disequilibrium (LD), i.e. non-independence across sites. Importantly, LD can accelerate mutational meltdown [14]. Empirically, LD has been detected even among unlinked sites [34,35]. Indeed, when *Ud* > 1, more LD is expected from background selection among unlinked sites than from background selection among linked sites in this regime [36].

The evidence for *U*_*d*_ > 1 is clearest for humans. A lower bound was derived [37] by assuming that mutations are deleterious only in the 55% of the 6×10^9^ base pairs in a diploid genome that are not dominated by dead transposable elements shared with chimpanzees, and that this 55% of the genome evolves 5.7% slower due to this constraint [38], implying that 5.7% of mutations were deleterious enough to be purged by natural selection. A conservative point mutation rate of 1.16×10^−8^ per base pair per replication [39] then yields *U*_*d*_ > 0.55×6×10^9^×0.057×1.16×10^−8^ = 2.2 per replication. This estimate is conservatively low, for five reasons. First, some mutations to transposable element regions are deleterious. Second, 1.16×10^−8^ is the lower bound of a 95% confidence interval on the point mutation rate [39]. Third, non-point mutations and beneficial mutations are neglected. Fourth, the constraint estimate [38] came from reference genomes that include some derived polymorphisms, which are less constrained than fixed differences. Fifth, under a distribution of fitness effects (DFE), some weakly deleterious mutations fix, and will not be included within the 5.7%. Some therefore argue that the human deleterious mutation rate is as high as *U*_*d*_ = 10 [40,41]. Humans have only a modestly higher point mutation rate than most other eukaryotes [39], making it likely that *U*_*d*_ >1 is common.

Capturing unlinked LD arising from *U*_*d*_ >1 requires simulating the entire genome rather than just a genomic region. This is computationally infeasible for most forward-time simulators (e.g., SLiM [42]), because the number of segregating mutations to be tracked scales with *U*_*d*_*N*, which becomes too large. Here we conduct whole genome simulations using a recently developed approach [26] that assumes that recombination occurs only at hotspots; instead of tracking every segregating mutation, we track only a summary of all mutations to date within each “linkage block” (between hotspots).

Here we ask how important mutational drought is relative to mutational meltdown, as a function of three main factors: 1) *U*_*d*_/*U*_*b*_, 2) the rate of environmental change, if any, and 3) the presence or absence of LD. We also confirm that the effect of LD on fitness fluxes and *N*_*crit*_ corresponds to that on *N*_*e*_ as assessed the usual way as the expected coalescence time of a neutral allele.

## Methods

### 1. Without linkage disequilibrium (LD)

#### Fixation probability

Independent evolution of each site (i.e. no LD nor epistasis) enables an analytical, one locus approach. Each copy of a mutation co-dominantly affects fitness by a factor of (1 + *s*), i.e. by (1 + *s*)^2^ for a homozygous mutant; this convention facilitates comparisons with simulations. The exact probability of fixation *p*_*fix*_(*s*) of a new mutation under the Moran birth-death process is then 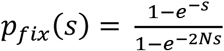. We take this from equation 1 of [43], adjusted for a diploid population of size *N*. It is exact for Malthusian fitness; per-generation fitness yields the same when an approximation is used [44].

#### Deleterious fitness flux

The best-studied DFE for deleterious mutations was inferred from the human site frequency spectrum of non-synonymous mutations (Table 2, row 9 of [45]). It fits a gamma distribution 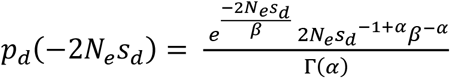, where Γ(*α*) is the gamma function, *α* = 0.169 is the shape parameter, and *β* = 1327.4 is the scale parameter. We divide by 2*N*_*e*_ = 23,646, yielding a DFE for non-synonymous *s*_*d*_ with α = 0.169, β = 0.056.

Many deleterious mutations occur in non-coding regulatory regions such as enhancers and promoters [46], and may have smaller mean deleterious effect size 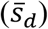 [47]. Structural mutations might have larger 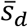. Departures from co-dominance affect the value of *s* corresponding to a given value of *p*_*fix*_(*s*). We therefore explored the sensitivity of our model to the deleterious DFE in two ways: by varying 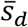 (adjusting the scale parameter while keeping the shape parameter fixed), and by varying the coefficient of variation (adjusting both the shape and scale parameters simultaneously to maintain a constant 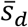).

The deleterious fitness flux (*v*_*d*_) is equal to the per-generation rate of new mutations *U*_*d*_*N*, times their fixation probability *p*_*fix*_(*s*_*d*_), times their homozygous impact on fitness 2*s*_*d*_ once fixed: 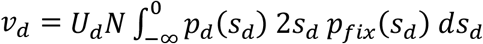. While selective effects 2*s*_*d*_ < −1 (worse than lethal) form part of the DFE we model, in practice they behave indistinguishably from lethal mutations.

#### Beneficial fitness flux

We follow others (e.g. [24]) to assume an exponential DFE with mean 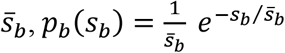 to obtain 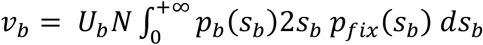.

#### Critical population size

We calculated the deleterious and beneficial fitness fluxes numerically in Mathematica 14.2.3 (Supplementary Notebook), using the DFEs *p*_*d*_(−2*N*_*e*_*s*_*d*_) and *p*_*b*_(*s*_*b*_), described above. The environmental change rate *δ*_e*n*v_ is either zero or in the range *δ*_e*n*v_= −10^−6^ to −10^−3.5^. We sometimes express the environmental change rate, for ease in interpretation, as the number of generations over which environmental change reduces fitness by 10%, with −0.1/*δ*_e*n*v_ = 10^5^ to 10^2.5^ generations. We consider even the fastest of these environmental change rates low because deleterious fixations cause more fitness loss than environmental change does. Coevolution, especially with parasites, likely makes the biggest contribution to *δ*_e*n*v_ [48].

We solve for *N*_*crit*_ as the value of *N* for which *v*_*n*et_ = v_b_ + *v*_*d*_ + *δ*_e*n*v_ = 0, and calculate the drought : meltdown ratio as 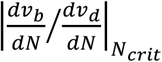. With linkage equilibrium, *N*_e_ = N.

The beneficial mutation rate and effect sizes will be larger in worse-adapted populations, but defining a “worse adapted population” is hard within a standard relative fitness formulation of population genetics. Higher *U*_*d*_ and *δ*_e*n*v_ would make populations worse adapted if all else were held equal, but in our modelling framework, these factors are offset by our consideration of perturbation to a higher *N*_*crit*_. Future work on less conventional models of absolute fitness under density-regulation (e.g., [49,50]) would be needed to explore the exact balance between these opposing considerations. Until then, the fact that there is a balance helps justify our use of the same values of *U*_*b*_ and 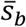 as we vary parameters such as *U*_*d*_ and *δ*_e*n*v_.

### 2. With LD

#### Simulation setup

Our forward time simulation method is described in [26]. Briefly, each genome consists of two haplotypes, divided into 23 chromosomes, each further segmented into *L* = 100 non-recombining ‘linkage blocks’ (which previous work found to be more than sufficient [26]). To reduce computational load, the fitness impacts of all mutations *i* within a given allele of linkage block *j* are not tracked separately, but only via their product *l*_*j*_ = ∏_*i*_ (1 + *s*_*i*_). Recombination occurs only at hotspots between adjacent linkage blocks. We simulated two recombination events per chromosome per meiosis [51].

In each time step of our Moran model, one of the *N* individuals is chosen uniformly at random to die. To replace it, we select two hermaphroditic parents with probability proportional to their fitness 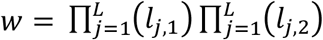, calculated across both haplotypes.

We sampled the numbers of new deleterious and beneficial mutations from Poisson distributions with means *U*_*d*_ and *U*_*b*_, respectively. We sampled selection coefficients from the same DFEs described in the previous section.

Simulations ran for 100N generations. Following [26], we ended a burn-in phase away from a clonal initial population 100 generations after a linear regression of the variance in fitness over the last 200 generations produced a slope less than 0.07/N. Net fitness flux *v*_*n*et_ was estimated post burn-in as the slope 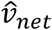 of log mean population fitness over time, without decomposition into *v*_*d*_ and *v*_*b*_.

#### Critical population size

*N*_*crit*_ is defined as the value of *N* for which *v*_*n*et_ = 0, but 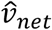 estimation is too noisy to find *N*_*crit*_ using purely deterministic root-finding methods, especially for small values of *N*_*crit*_. To account for this stochasticity, we performed 10 independent replicates of the following procedure for each parameter set. We first bracketed *N*_*crit*_, with *N* = 1500 and 6000 as initial guesses, progressing outward by a factor of two as needed to obtain 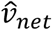 values both above and below zero. Then we iterated the secant method until 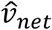 changed by < 15%. Then we fitted a straight line to all evaluated {N, 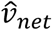} data points for which *N* was within a factor of 3 from the final evaluated value. We estimated *N*_*crit*_ for each replicate as the zero-intercept of this fitted line, and used the mean of these estimates across the 10 replicates for subsequent simulations of matched *N*.

#### Drought : meltdown ratio

We estimated *v*_*d*_ and *v*_*b*_ from the set of mutations whose time of final fixation lay between the end of the burn-in phase and the end of the run. Tracking fixations does not exactly track mean population fitness due to fluctuations caused by polymorphisms. For computational efficiency, the linkage block method described so far makes information about individual fixed mutations inaccessible. In simulations intended to estimate the drought : meltdown ratio, we made them accessible via tree sequencing recording [52] tracking all non-neutral mutations, including their time of fixation. We calculated 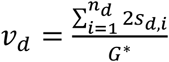 and 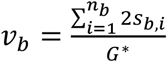, where *G*^∗^ is generations elapsed since burn-in ended, and *n*_d_ and *n*_*b*_ are the numbers of deleterious and beneficial mutations that fixed during this time.

We numerically estimated 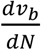 and 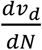 as 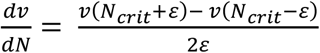, where we chose *ε* = 150 individuals as a balance between reducing noise and avoiding curvature in *v*. We did this by running simulations for *N*_*crit*_ + *ε* and N_cr*it*_ − *ε*.

## Results

Without LD, where *N*_*e*_ = *N*, Whitlock’s [24] choice of *U*_*d*_/*U*_*b*_ = 1000, combined with the deleterious DFE from [45], and a conservatively low 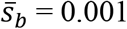, with both kinds of mutation co-dominant, yields *N*_*crit*_ ≃ 3666 (Figure 2A, solid blue and red lines intersecting at dotted red vertical line). An analytical approximation based on an exponential DFE (equation (9) in [24] scaled by 2 to account for the Moran model) underestimate 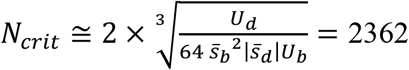. This discrepancy likely arises because our DFE from [45] has a coefficient of variance of 2.4, in contrast to 1 for an exponential distribution. Directly integration of Eq. 3 in [24], which comes prior to the exponential assumption, resolves the discrepancy, to produce *Ncrit* ≃ 3665. Figure S1 illustrates similar agreement across the range of *Ud*/*Ub* values. We note that *N*_*crit*_ values in [24] are intended to represent *N*_*e*_ not census *N*.

**Figure 2.**
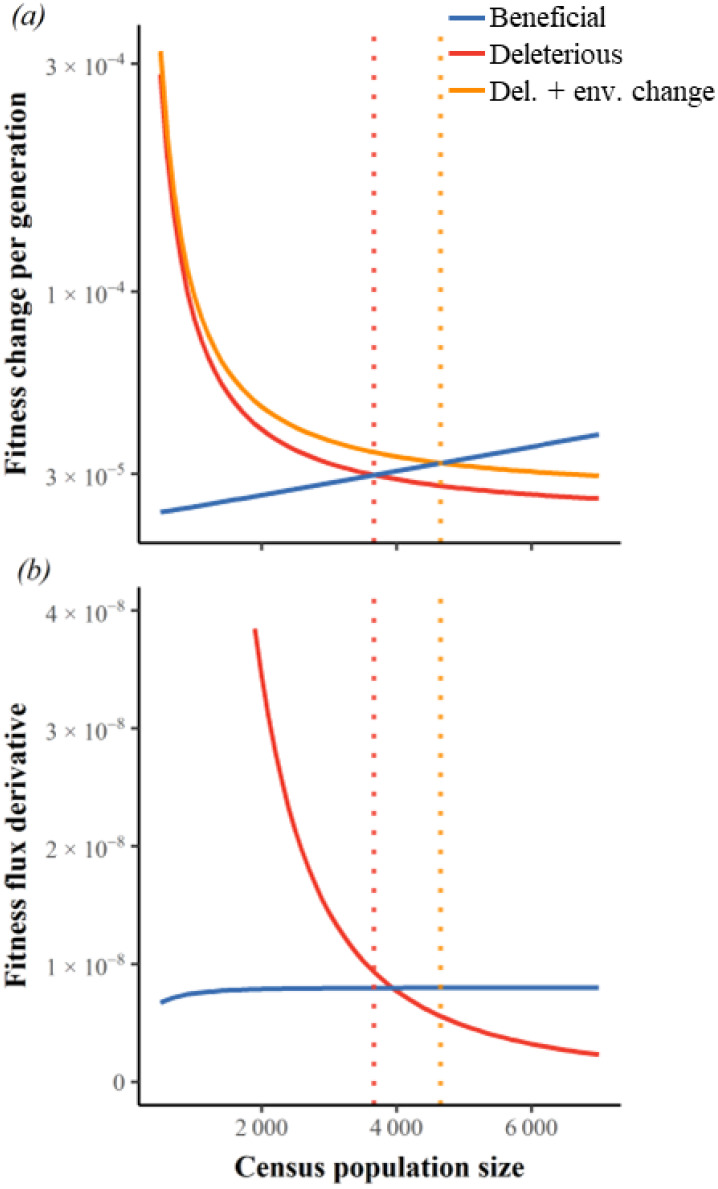
Extinction vortices are driven more by mutational drought than by mutational meltdown. Vertical lines indicate *N*_*crit*_, defined by the intersection in (a) of beneficial fitness flux (blue) with either deleterious fitness flux (red), or with deleterious flux plus environmental deterioration (orange). Environmental change adds a constant *δ*_*en*v_ = −1.5×10^−5^ per generation to deleterious flux in (a) and hence does not change its slope (red line in (b)). The ratio of the y-values in (b) at *N*_*crit*_ indicates the relative importance of mutational drought (blue) vs. mutational meltdown (red). *U*_*d*_ = 2, *U*_*d*_/*U*_*b*_ = 1000, 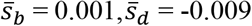 via the DFE from [45], and there is linkage equilibrium.

At *N*_*crit*_ in the absence of environmental change (dotted red vertical line in figure 2b), declining *N* has 85% of the impact on beneficial flux as it has on the deleterious flux that is normally assumed to be the sole cause of an extinction vortex according to mutational meltdown theory. When adaptation must also counter environmental change, larger populations enter the extinction vortex (figure 2a, orange vertical dotted line illustrates *N*_*crit*_ ≃ 4656 when environmental change reduces fitness by 10% over ∼6,000 generations). Even modest environmental change makes the impact of declining *N* on beneficial flux larger than that on deleterious flux (blue line is above red along dashed orange vertical line in figure 2b). In other words, environmental change makes mutational drought more important than mutational meltdown.

Indeed, the drought : meltdown ratio is > 1 with *U*_*d*_/*U*_*b*_ = 1000 when slow environmental change causes fitness to decline by 10% over as many as ∼20,000 generations (figure 3a, right, grey, dot-dashed vertical line). When *U*_*b*_ is 10 times higher, faster but still modest environmental change is needed to drive the drought : meltdown ratio > 1 (figure 3a, left, blue, dot-dashed vertical line, ∼4000 generations for fitness to decline by 10%). We interpret both these rates of environmental decline as rather slow because they correspond to only 0.19× and 0.28× the rates of fitness decline due to deleterious fixations for *U*_*d*_ = 2 (figure 3b, colour-matched intersections with vertical lines). Environmental change can drive the drought : meltdown ratio > 1 without excessively elevating *N*_*crit*_ (figure 3c steep rise in *N*_*crit*_ occurs well to the left of the color-matched vertical line).

**Figure 3.**
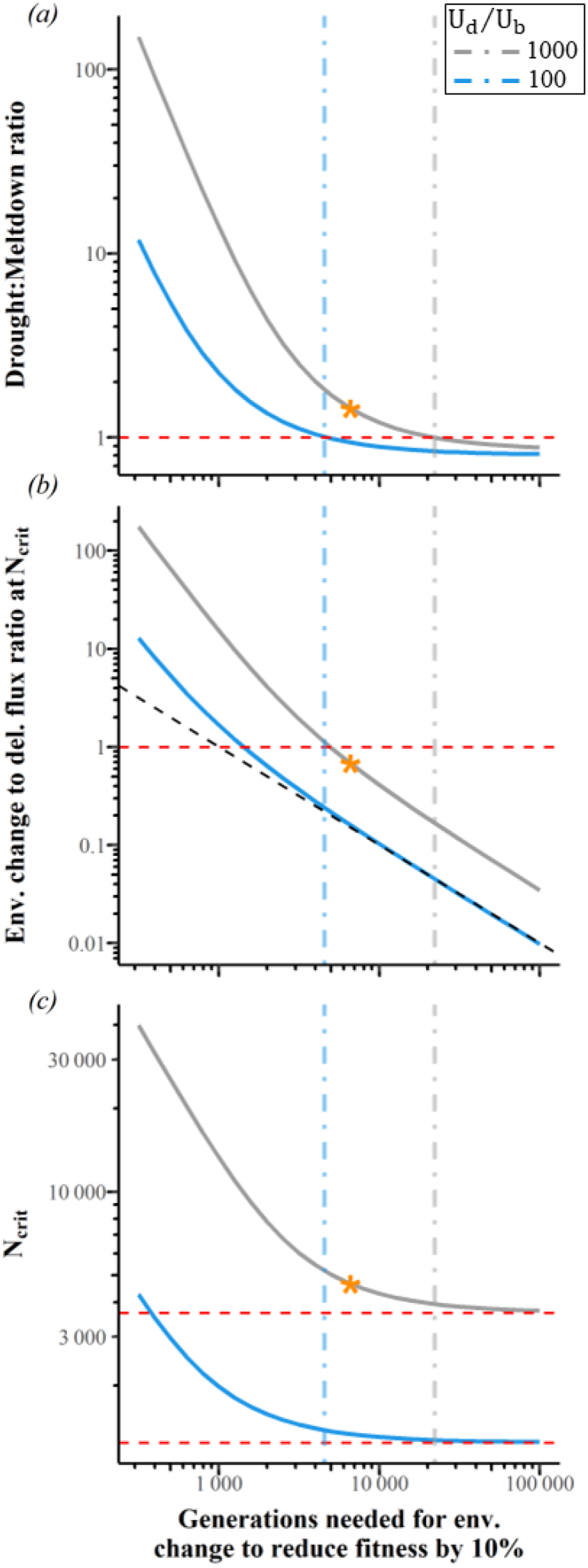
Slow environmental degradation is sufficient to make mutational drought more important than mutational meltdown. 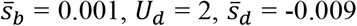 via the DFE from [45]. Linkage equilibrium is assumed. The orange star indicates the example shown in figure 2. Vertical lines represent where the drought : meltdown ratio = 1. (b) We consider environmental change to be relatively slow below the red dashed horizontal line, where mutational degradation makes a bigger contribution to fitness decline. Diagonal black dashed line in (b) represents proportionate change in the x-axis and y-axis; departure from this indicates increased deleterious flux in the face of rapid adaptation. Red dashed lines in (c) indicate asymptotes, i.e. no environmental change.

When beneficial mutations are scarce (higher *U*_*d*_/U_b_), mutational drought is relatively more important, with the drought : meltdown ratio saturated by the *U*_*d*_/*U*_*b*_∼1000 value (figure 4a) used in figure Lowering *U*_*b*_ has two opposing effects. First, it proportionately lowers the beneficial fitness flux *v*_*b*_ and its derivative 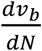 (i.e. it reduces the slope of the blue line in figures 2A and consequently reduces the height of the blue line in figure 2B). Second, lower *v*_*b*_ moves the intersection *N*_*crit*_ to the right, i.e. towards larger values of *N*, which lowers 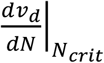 without substantially changing 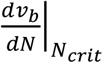 (figure 2B, red and blue lines, respectively). The flat drought : meltdown ratio to the right of figure 4a shows that when beneficial mutations are scarce at high *U*_*d*_/*U*_*b*_ ≥ 1000, these two effects cancel each other out.

**Figure 4.**
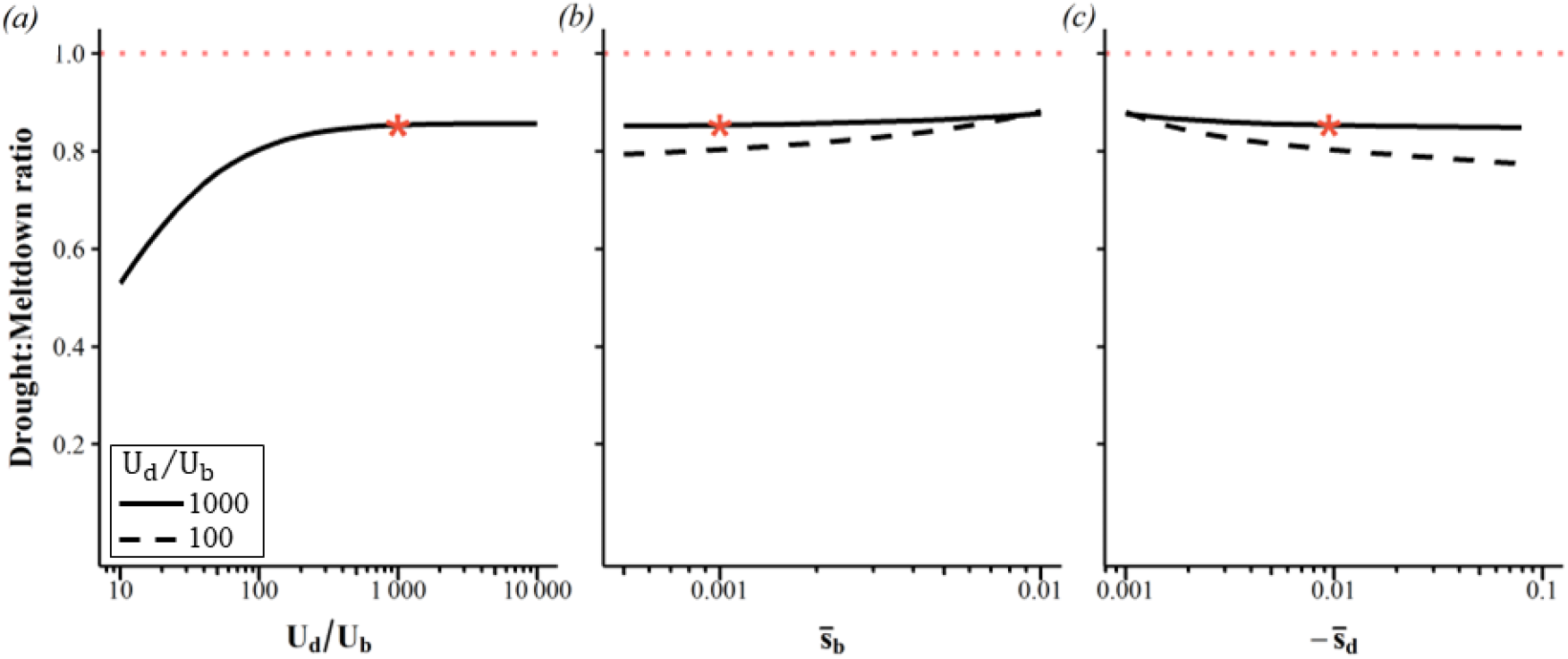
The drought : meltdown ratio is fairly insensitive to parameter value choices. Linkage equilibrium is assumed, with no environmental change. (a) Mutational drought becomes somewhat less important when (a) beneficial mutations are abundant, (b) beneficial effects are weak, and (c) deleterious effects are strong. Where not otherwise specified, 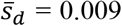 via the DFE from [45], 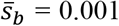. *U*_*d*_ = 2 throughout, but results for different *U*_*d*_ are superimposable. The red star indicates the example shown in figure 2 with no environmental change. See figure S2 for corresponding changes in *N*_*crit*_.

Weaker beneficial effects (figure 4b), and stronger deleterious effects (figure 4c) make mutational drought only slightly less important. These effects are only slightly larger when beneficial mutations are more common (figures 4b and 4c dashed curves). This makes our results insensitive to the difficulties of empirically inferring selective effect sizes and their dominance coefficients, and to the fact that global epistasis might cause selective effect sizes to vary across scenarios. Mutational drought is relatively more important when the deleterious DFE has a higher coefficient of variation, while holding 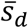 constant (Figure S3). The non-synonymous DFE we use from [45] is overdispersed, and mixing this with DFEs of different 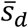 is likely to further increase the overdispersion.

With environmental change, dependence on *U*_*d*_ and *U*_*b*_ does not simplify to dependence on their ratio, as in figure 4a. When faster environmental change and more scarce beneficial mutations push the drought : meltdown ratio above 1 (figures 5a and 5b), lower *U*_*d*_ makes mutational drought more rather than less important (figure 5a). The drought : meltdown ratio only falls substantially below 1 under the slower environmental change scenario, and only when *U*_*d*_/U_b_ is small (figure 5a, left portion of dashed blue). This scenario entails exceptionally small *N*_*crit*_ (figure 5e). Under realistically high *U*_*d*_, high drought : meltdown ratios are found when beneficial mutations are scarce (figure 5b, solid lines to right), making *N*_*crit*_ high (figure 5f), and thus environmental change at *N*_*crit*_ is relatively fast compared to a slower rate of deleterious fixations (figure 5d). The drought : meltdown ratio is mostly insensitive to selection coefficients, although it rises for very small 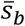 (figure S4).

**Figure 5.**
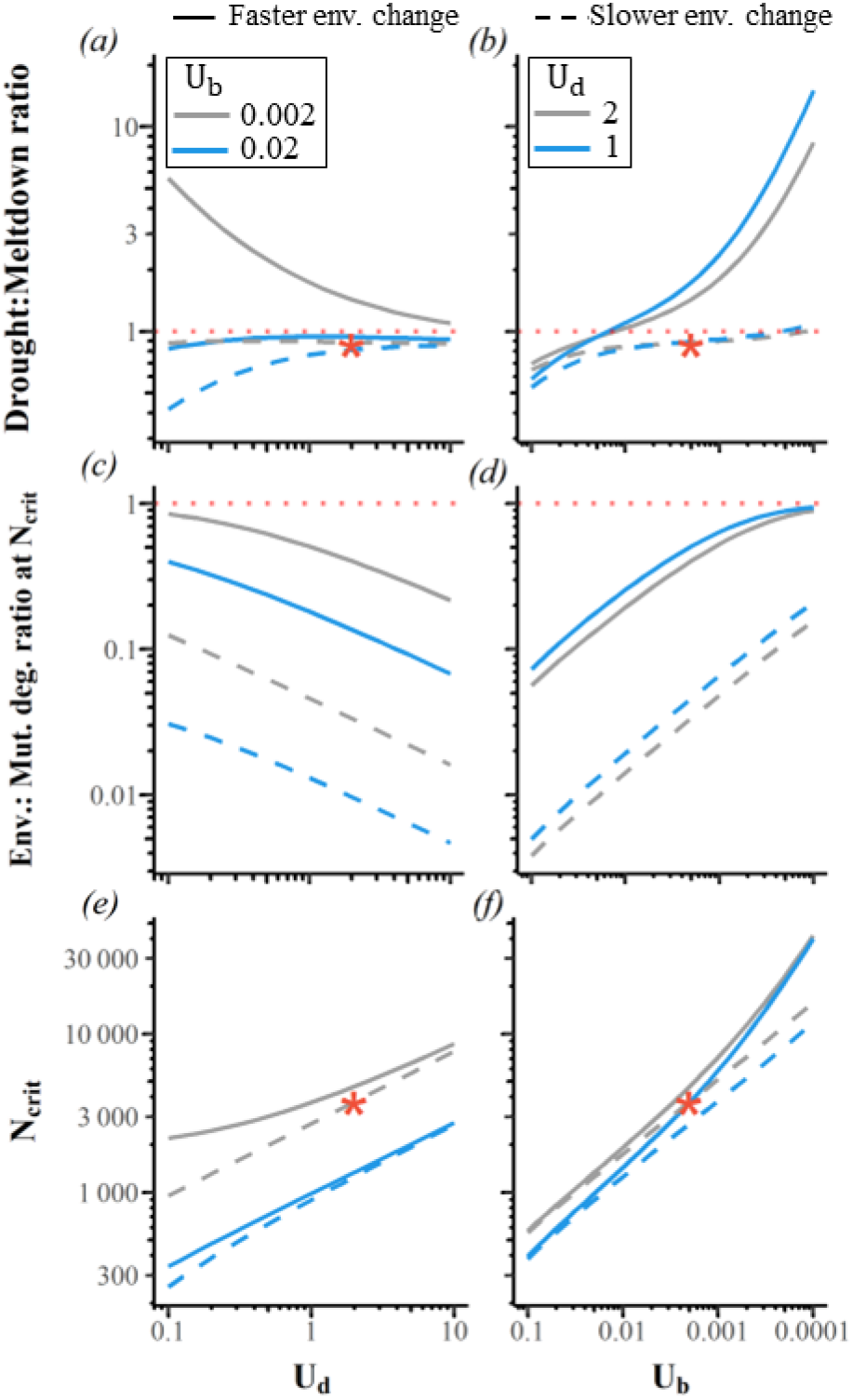
The drought : meltdown ratio only falls substantially below 1 for slower environmental change, and only when *U*_*d*_/*U*_*b*_ is small. (a) Faster environmental degradation, combined with low *U*_*b*_, can make mutational drought more rather than less important for lower *U*_*d*_. (a) and (d) can be compared to the constant environment in figure 4a, where only the ratio *U*_*d*_/U_b_ matters. Reverse scaling of the *U*_*b*_ x-axis of (d-f) is used to facilitate comparison of (d) with (a) and with figure 4a. In our “fast” environmental change scenario (solid lines; *δ*_*en*v_= −1.5×10^−5^ per generation, matching the example in figure 2), fitness loss from environmental degradation is comparable to, though still slower than, fitness loss from deleterious fixations (y-axis values modestly <1 in (c-d)). In our “slow” scenario (dashed lines; *δ*_*en*v_ = −10^−6^), environmental degradation is much slower than mutational degradation (c-d), making results comparable to the constant environment example from figure 2 (red star). Independent sites model with 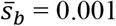 and 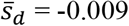 via the DFE from [45].

The presence of LD, which we capture in simulations with *U*_*d*_ = 2 and no environmental change, produces a modest 8% increase in the drought : meltdown ratio, from 0.85 to 0.92, as beneficial flux becomes slightly more sensitive to *N* (figure 6d, blue points are above blue line). This goes away when we quantitatively reduce LD by increasing the number of chromosomes from 23 to 50 (figure S4a).

**Figure 6.**
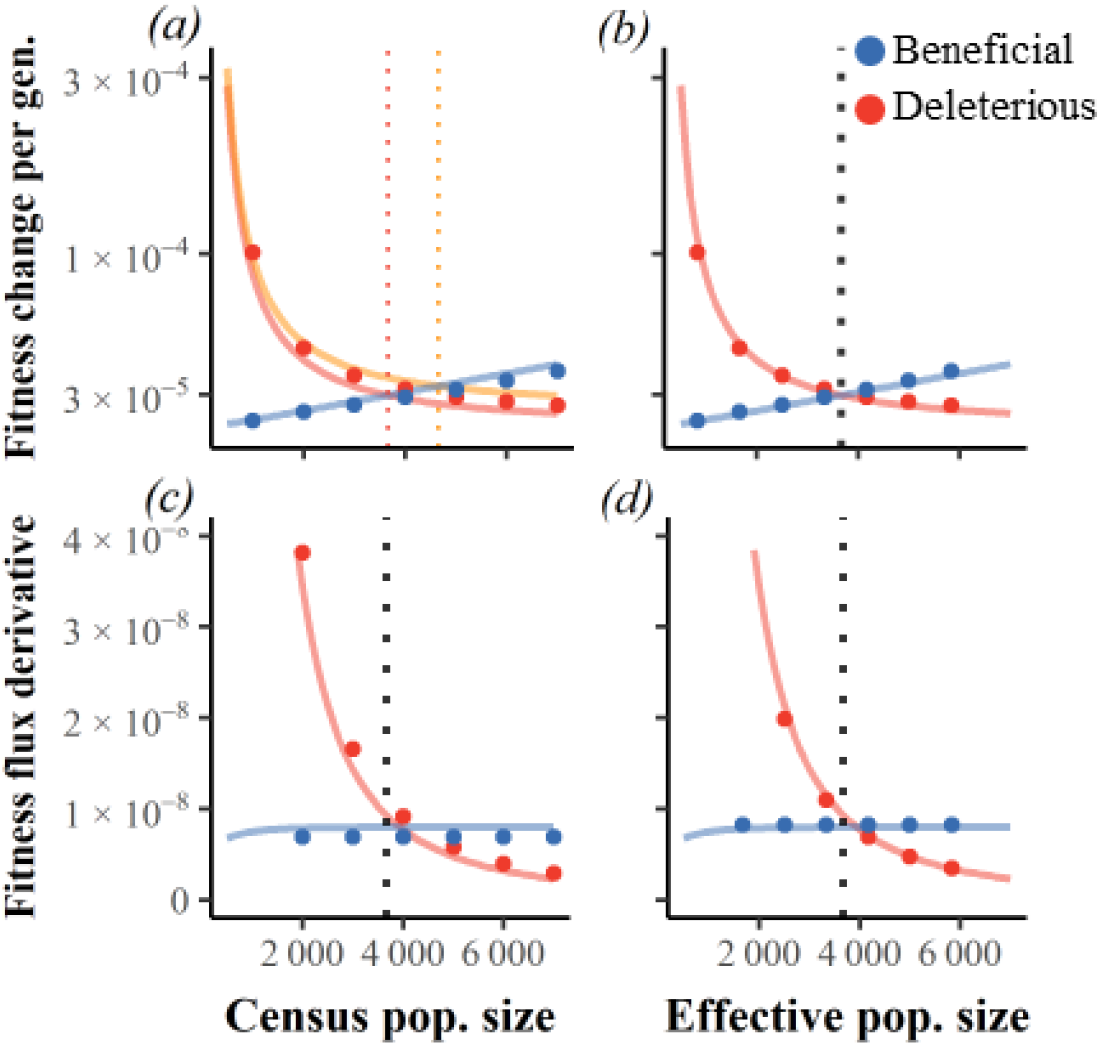
Fitness fluxes scale with effective population size under linkage disequilibrium. Continuous red and blue lines are analytical results given linkage equilibrium, while dots are simulation results with LD. (a) and (b) add simulation results to the static environment example of figure 2. In (c) and (d), the x-axis for simulation results is scaled such that intersection occurs at the same “effective” *N*_*crit*_ as found without LD. Linkage disequilibrium makes mutational drought slightly more important. *U*_*d*_ = 2, *U*_*d*_/U_b_ = 1000, 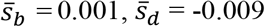 via the DFE from [45].

LD increases *N*_*crit*_ from 3666 to 4613 (figure 6a, solid dots), in a way that is qualitatively robust to adding more chromosomes (figure S4b). The *N* = 4613 population with LD can thus been seen to have *N* = 3666, such that 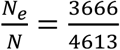. Beyond *N*_critt_, rescaling describes *v*_*d*_ and *v*_*b*_ well, as seen by the close match between analytical lines and simulated points when the x-axes of figures 6a and 6c are rescaled according to 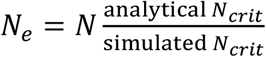 to produce figures 6b and 6d. LD slightly increases the magnitude of fitness fluxes at their *v*_*d*_ = *v*_*b*_ intersection (figure 6b).

The reduction in *N*_*e*_ could be due to beneficial mutations interfering with each other (clonal interference decreasing beneficial flux), and/or hitchhiking (beneficial mutations increasing deleterious flux), and/or background selection (deleterious mutations both decreasing beneficial flux and increasing deleterious flux). We assessed *N*_*e*_ in control simulations with *U*_*b*_ = 0, while using matching values of *N* = *N*_*crit*_ in the corresponding scenario with *U*_*b*_ > 0. Equal values of coalescent *N*_*e*_/N in these controls (figure S6, red vs blue) points to background selection as the cause. Assessing *N*_*e*_ via the reduction in fitness fluxes (figure S6, black circles) yields similar values of *N*_*e*_/N.

## Discussion

Mutational meltdown describes an extinction vortex in which small populations cannot purge slightly deleterious mutations, whose fixations handicap the population, making it still smaller. Here, we find that a shortage of beneficial mutations appearing in smaller populations creates mutational drought, a distinct kind of extinction vortex. Mutational drought is only slightly less significant than mutational meltdown in the absence of environmental change, and becomes more important in the more realistic scenario of even a slowly changing environment. Mutational drought is modestly more significant with LD, and more significant when beneficial mutations are rare. Beneficial mutations are thus either common, or else their population-size-dependent scarcity is critical – in neither case can population genetic models ignore beneficial mutations. The relative roles of drought and meltdown are remarkably insensitive to selection coefficients, and only modestly dependent on the relatively frequencies of beneficial vs. deleterious mutations. The greatest parameter value sensitivity is to the rate of environmental change, which if fast, can drive a huge excess of drought over meltdown.

We have ignored population structure. In structured populations, the synergistic interaction between mutation accumulation and demography makes fragmented metapopulations more vulnerable to extinction than panmictic populations of equivalent size [53]. However, fragmentation might not affect mutational drought the same way. Indeed, in asexual populations, fragmentation has been empirically found to improve adaptation given ruggedness of the fitness landscape [54]. When environments vary among demes, high dispersal can trigger a ‘migrational meltdown’ by swamping local adaptation [55]. On the other hand, under hard selection, high migration rates are required to buffer demes against local mutational meltdown [56]. It is not yet known what migration rates are needed to similarly buffer demes against local mutational drought.

The role of beneficial mutations in population persistence studied here plays out over much longer timescales than processes of “evolutionary rescue” in previous population genetic (discrete) models, although in line with the broader use of the same term in the context of quantitative genetics models (see [28]). In the broad sense, evolutionary rescue refers to scenarios where genetic adaptation struggles to keep up with ongoing environmental change, over an indefinitely long period of time [57–61]. Within the narrow sense, population genetic models of evolutionary rescue consider a population whose birth rate is initially less than its death rate, e.g. following a single abrupt environmental change that can be remedied by one or several beneficial mutation(s) [62–67]. Here we used a population genetic model to consider evolutionary rescue in the broad sense more commonly treated by quantitative genetic models, across an indefinitely long series of beneficial and deleterious fixations.

Conservation management and species recovery programs assess the ‘minimum viable population’ and the factors influencing it [68–71]. We follow [72] in defining *N*_*crit*_ as the lowest population size that avoids an extinction vortex. This is a different kind of minimum viable population to the usually considered empirical predictor of probability of persistence after a specified period of time, e.g. 1000 years [71,73]. Our *N*_*crit*_ concerns the ability to resupply the population with new beneficial mutations to balance both the accumulation of deleterious mutations, and ongoing environmental change, to achieve long-term viability. *N*_*crit*_ is the gateway to the extinction vortex below which some combination of mutational meltdown and mutational drought contribute. Our *N*_*crit*_ is relevant over the long timescales of multiple sweeps, making our results applicable more clearly to species than to populations, reliant on mutation rather than migration as the source of new variants.

The assessed extinction risk of vertebrate species in the Anthropocene [74] includes assessed long-term risks to environmental conditions [75]. Current conservation efforts often focus on emergency revitalization programs with tools that might only achieve short-term persistence (e.g., [76]), potentially creating trade-offs between long-term and short-term objectives [77]. Short-term goals may be problematically prioritized under triage from restricted budgets [78,79] over achieving long-term sustainable populations [80,81].

Some conservation geneticists advocate genetic rescue, whereby gene flow from a few introduced individuals helps small fragmented populations [82–84]. The genetic rescue paradigm aims to maximize genetic diversity to reduce the risk of extinction [82,85,86]. However, gene flow from large, outbred populations may introduce too many recessive deleterious alleles to be easily purged, with deleterious effects that might quickly have devastating consequences within a small inbred population [6,87,88]. Abandoning the genetic rescue paradigm might limit the risk of deleterious accumulation in the short term. Our results on mutational drought help emphasize that this might still be a mistake over longer timescales if a shortage of adaptive mutations later poses an extinction risk. Fortunately, according to simulations, the benefits of genetic rescue can be obtained while minimizing harm by steadily introducing migrants from other small inbred populations from which recessive deleterious mutations are already purged [6,88], as has occurred naturally in Florida scrub-jays [89].

## Acknowledgements

We thank Daniel Smith, Ryan Gutenkunst, Elise Lauterbur, and David Enard for their thoughtful comments on parts of this study, and Naman Panday for sharing his git expertise. We also thank Jerome Kelleher, Peter Ralph, and Yan Wong for helpful discussions about tree sequence recording. Finally, we extend our thanks to the anonymous reviewers, whose comments which have greatly improved this work.

## Data Availability

Analytical results conducted in Wolfram Mathematica. Simulation code was written in C, and graphs produced with R. Simulation code and scripts have been archived at [90].

