## Supplementary Figures for "Extinction vortices are driven more by a shortage of beneficial mutations than by deleterious mutation accumulation"

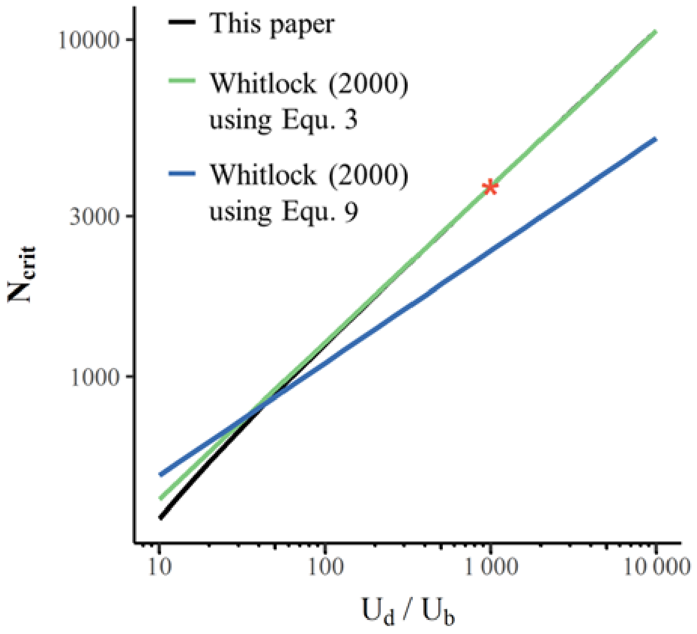


**Figure S1. Our analytical treatment of the critical population size is better approximated by Whitlock's (2000) Equation (3) than by his Equation (9).** To use Equation (3), we take $v_d$ from there rather than from the approximate Equation (4) to produce an alternative to Equation (9). Whitlock's predictions were multiplied by 2 to account for the higher genetic drift of the Moran process relative to the Wright-Fisher model. Results are shown for $U_d = 2$ and varying $U_b$, but we confirmed that results varying $U_d$ instead, while holding $U_b$ =0.002, are superimposable. $\bar{s}_d$ = -0.009 via the DFE from Kim et al. (2017) from their Table 2 (row 9) and $\bar{s}_b$ = 0.001. Red star represents our example of $U_d/U_b$=1000 in main figures 2 and 6 with no environmental change.

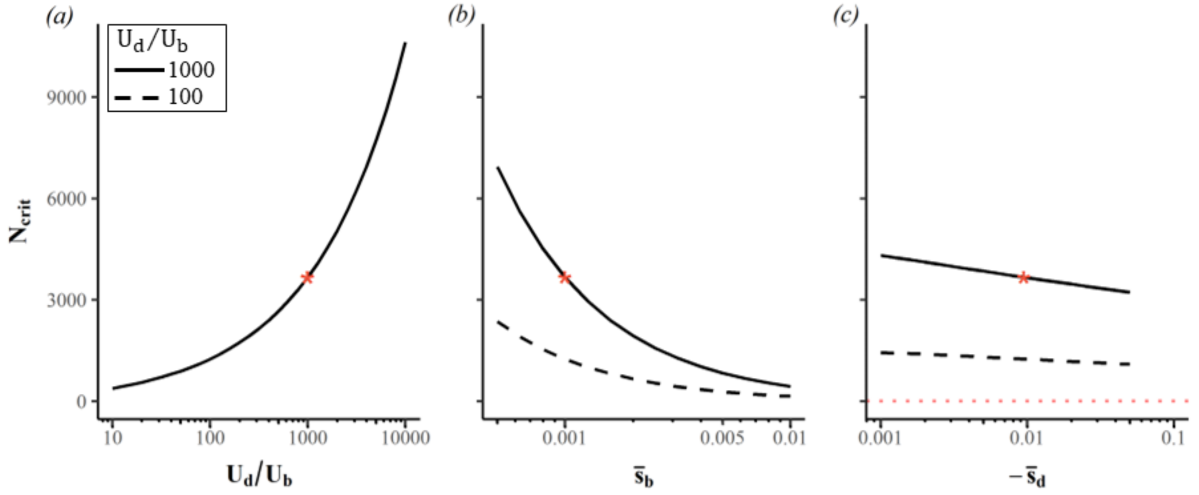


**Figure S2**. **$N_{crit}$ dependence on parameter value choices.** Linkage equilibrium is assumed, with no environmental change. (a) $N_{crit}$ is much lower when beneficial mutations are abundant (low $U_d/U_b$). $N_{crit}$ drops less with (b) stronger beneficial effects and less again with (c) stronger deleterious effects. Panels (a) and (b) use $\bar{s}_d$ = 0.009 via the DFE from Kim et al. (2017). Panels (a) and (c) use $\bar{s}_b$ = 0.001. $U_d$ = 2 throughout, but results for different $U_d$ are superimposable. The red star indicates the example shown in Figure 2 with no environmental change.

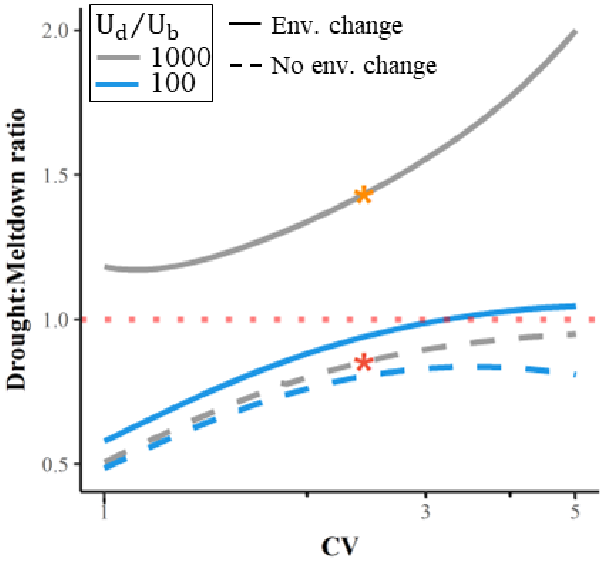


**Figure S3**. **Drought becomes more important relative to meltdown when the deleterious DFE is overdispersed.** Linkage equilibrium is assumed. To isolate the effect of the coefficient of variance (CV), we hold the mean deleterious effect size constant at the $\bar{s}_d$ corresponding to the DFE from Kim et al. (2017), and adjust the parameters of the gamma distribution. Beneficial mutations have an exponential distribution with $\bar{s}_b$ = 0.001. The red and orange stars indicate the two scenarios in Figure 2, with no environmental change and with environmental change ($\delta_{env}$= -$1.5\times10^{-5}$ per generation), respectively.

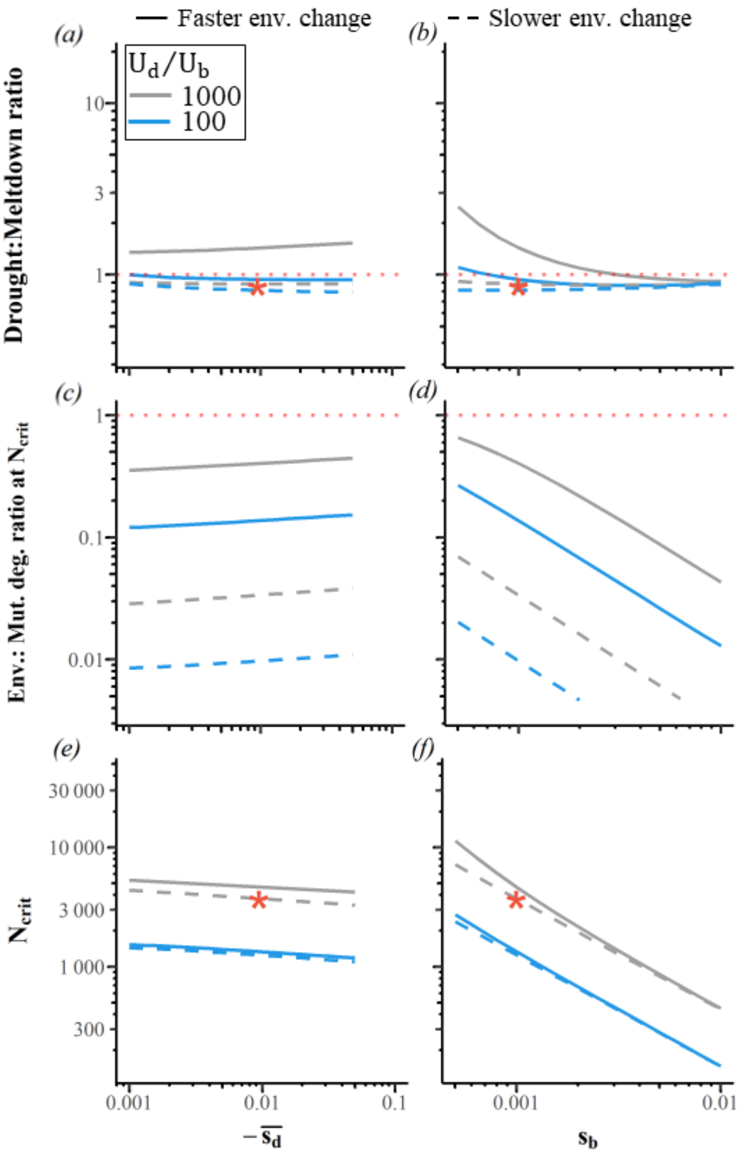


**Figure S4. The drought : meltdown ratio is largely insensitive to selection coefficients, but rises for small $\bar{s}_b$.** As in Figure 5, the red star indicates the example shown in Figure 2 with no environmental change. "Faster env. change" (blue lines) represents a fitness reduction of $\delta_{env}$= -1.5×10$^{-5}$ per generation, and "slower env. change" (black lines) represents $\delta_{env}$ = -10$^{-6}$. Results are shown for $U_d = 2$ and varying $U_b$, but results varying $U_d$ instead, while holding $U_b$ =0.002, are qualitatively similar. The y-axes scales match those in Figure 5, to facilitate comparison.

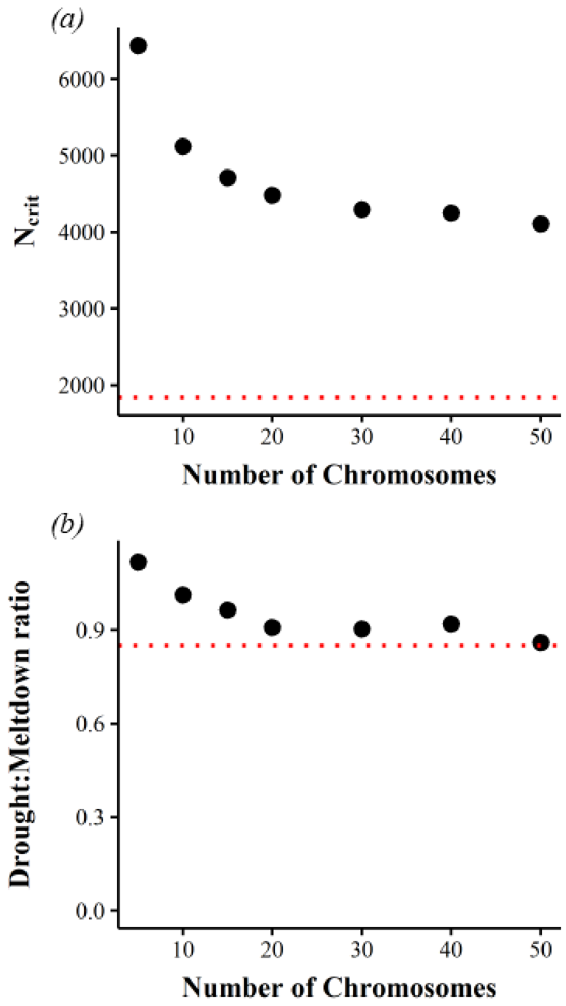


**Figure S5**. **More recombination does not make $N_{crit}$ converge to the expectation for independent loci (a), but this does not affect the drought : meltdown ratio (b).** $\bar{s}_b$ = 0.001, $U_b$ = 0.002, $U_d$ = 2, $\bar{s}_d$ = -0.009 via the DFE from Kim et al. (2017), $L$ = 100 per chromosome. Red dashed lines show analytical expectations in the absence of LD.

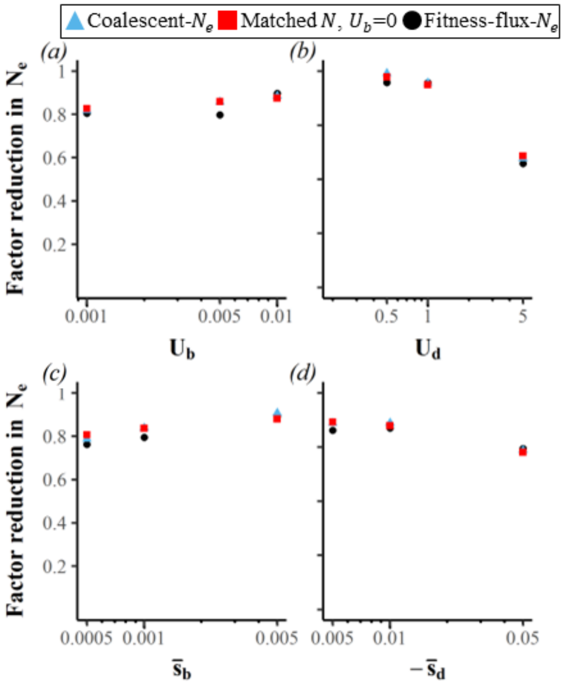


**Figure S6**. **The reduction in effective population size below census size is driven by background selection.** The reduction derived from fitness fluxes $\frac{\text{analytical } N_{crit}}{\text{simulated } N_{crit}}$ (black circles) is close to $N_e/N$ from coalescence time (blue triangles). Both align with coalescent $N_e/N$ derived from simulations of matched census $N$ with deleterious mutations alone ($U_b$ = 0, red squares). Where not otherwise specified, $\bar{s}_b$ = 0.001, $U_d$ = 2, $U_b$ = 0.02, and $\bar{s}_d$ = -0.009 via the DFE from Kim et al. (2017). There is no environmental change.
